# *Mus Musculus* papillomavirus MmuPV1 resists restriction by human APOBEC3B

**DOI:** 10.1101/2025.11.10.687577

**Authors:** Xingyu Liu, Andrea Bilger, Denis Lee, Prokopios P. Argyris, Jiarui Chen, Ella Ward-Shaw, Emilia Barreto Duran, Yu-Hsiu T. Lin, Cameron Durfee, Sang Hyun Chun, Mahmoud Ibrahim, Joshua Proehl, Allen J. York, Paul F. Lambert, Reuben S. Harris

## Abstract

The single-stranded DNA deaminase APOBEC3B (A3B) is capable of potently restricting the replication of a range of viruses including retroviruses (cDNA) and herpesviruses (genomic DNA). However, these and likely other DNA virus families have evolved host species-specific counter-defenses that are equally potent and serve to protect viral DNA from restriction. Although high-risk human papillomavirus (HPV) infection triggers A3B upregulation, potentially as part of an antiviral response, the impact of this restriction factor on papillomavirus replication and pathogenesis has yet to be assessed. To study human A3B antiviral function in the absence of a species-specific counter-defense, here we ask whether human A3B is capable of restricting *Mus musculus* papillomavirus (MmuPV1) *in cellulo* and *in vivo*. First, we created human A3B and catalytic mutant A3B-E255A expressing FVB/N mice. Second, MmuPV1 gene expression and replication was quantified in primary keratinocytes from these animals and, surprisingly, enzymatically active human A3B caused no measurable impairment in viral transcript or DNA accumulation. Third, A3B, catalytic mutant A3B-E255A, and nontransgenic FVB/N animals were infected with MmuPV1 and similar pathologies were found regardless of A3B functionality. Thus, despite likely never being exposed to human A3B during evolution, MmuPV1 appears to be unaffected by this potent, primate-specific antiviral factor. These results suggest that MmuPV1 and perhaps papillomaviruses more broadly possess a conserved mechanism to efficiently escape restriction by human A3B and related DNA deaminases.

**IMPORTANCE:** Human papillomaviruses (HPVs) are nearly ubiquitous, and persistent infection with high-risk types causes approximately 5% of cancers worldwide. Although HPV vaccination is effective for preventing infection, insufficient global coverage and a rising incidence of HPV-associated malignancies, such as oropharyngeal carcinoma, highlight the need to understand innate virus clearance mechanisms. APOBEC3 enzymes are a central component of the mammalian innate immune system and are hypothesized to restrict papillomavirus infection, particularly between species. Here, we establish mice that express the human antiviral enzyme APOBEC3B (A3B). Surprisingly, we find that human A3B is incapable of blocking the replication of a murine papillomavirus (*Mus musculus* papillomavirus 1, MmuPV1) in relevant primary cells from these animals or in infected tissues *in vivo*. These findings highlight the complexity of teasing apart host-pathogen interactions and suggest that papillomaviruses may have a general mechanism for escaping restriction by antiviral enzymes such as A3B.

## INTRODUCTION

Papillomaviruses (PVs) are small double-stranded DNA viruses and one of the most prevalent sexually transmitted pathogens worldwide (1–4). Human papillomavirus (HPV) infects the basal keratinocytes of epithelial tissues of the oral and genital regions of the body. HPV infections are often asymptomatic and cleared rapidly. However, a subset of infections can become persistent and long-lasting. Persistent high-risk HPV infections are critical drivers of tumorigenesis, with more than 95% of cervical cancers and roughly 30% of head and neck cancers driven by these virus types (5, 6). Understanding viral clearance mechanisms is therefore an attractive strategy for developing future therapies that help to reduce the risk of HPV-associated malignancies.

A major component of the host defense against DNA virus infection is the APOBEC family of enzymes that catalyze the deamination of cytosine to uracil in single-stranded DNA (7–9). In response to infection, APOBECs are often upregulated and capable of introducing mutations that compromise viral genomes and lead to abortive infections. APOBEC mutational signatures are frequently observed in HPV genomes and are also among the predominant mutation sources in HPV-related cancers (10–14). Our previous work showed that, among the APOBEC family members, APOBEC3B (A3B) is uniquely upregulated during high-risk HPV persistent infection in human immortalized primary keratinocytes (15). However, despite mutational footprints on viral genomes, experimental evidence suggests that A3B may be incapable of efficiently restricting HPV infection in human keratinocytes (16, 17). These observations raise the possibility that HPV has developed strategies to tolerate or evade A3B restriction activity.

*Mus musculus* papillomavirus type 1 (MmuPV1) is an emerging murine papillomavirus that can disseminate between laboratory mice in natural infection (18–20). Since its isolation in 2011, MmuPV1 has become a powerful experimental system to study papillomavirus infection and pathogenesis *in vivo* in murine models (18, 19, 21). The fact that MmuPV1 replicates *in vivo* suggests that it may have evolved mechanisms to counteract host innate immune barriers including the single murine APOBEC3 enzyme. Moreover, given the fact that mice naturally lack a direct ortholog of human A3B due to its existence exclusively in primates, there has been no co-evolutionary adaptation between MmuPV1 and this potent antiviral enzyme. Thus, MmuPV1 would be expected to be highly susceptible to restriction by human A3B.

In studies here, we have generated a novel transgenic mouse model that conditionally expresses human A3B to directly test whether MmuPV1 can be restricted by this antiviral factor. Using both *in vitro* and *in vivo* systems, we quantified viral transcription and replication in the presence and absence of human A3B. Our findings demonstrate that MmuPV1 is resistant to A3B-mediated restriction, providing new insights into papillomavirus-APOBEC interactions and suggesting the existence of a conserved counteraction mechanism capable of functioning across species.

## MATERIALS AND METHODS

### Mice

FVB/N *Rosa26::CAG-LSL-A3Bi* and FVB/N.*Rosa26::CAG-LSL-A3Bi-E255A* knock-in mice were generated at the Gene Targeting & Transgenic Facility at the HHMI Janelia Campus using Cas9 cleavage and homologous integration of reported minigene constructs (22). Briefly, the human A3B minigene construct, sgRNA targeting *Rosa26* (5’-CTCCAGTCTTTCTAGAAGAT-3’), and Cas9 protein (ThermoFisher Scientific, A36498) were co-injected into one-cell embryos in FVB/NCrl background (Charles River Laboratories, Strain: 207). Individual founders were identified by PCR and bred to identify the subset capable of germline transmission of the A3B minigene. *β-actin::Cre* mice were purchased from Jackson laboratory (FVB/N-*Tmem163^Tg(ACTB-cre)2Mrt^*/J; Strain #:003376). The transcription stop cassettes (*loxP-STOP-loxP*, LSL) were removed by crossing *Rosa26::CAG-LSL-A3Bi* and *Rosa26::CAG-LSL-A3Bi-E255A* to *β-actin::Cre* animals. The resulting offspring express active human A3B enzyme or the catalytic mutant A3B-E255A protein in nearly all tissues at similar levels and, apart from the E255A substitution, are otherwise isogenic. Primers are listed in **Table S1**.

### Virus preparation

Virus stocks were prepared as described (23). Warts were collected from ear, skin, tail, anus, and other infected areas of nude mice and soaked in PBS overnight at 4°C. Then warts were homogenized by PowerGen 125 (ThermoFisher Scientific, PG125). The lysates were incubated with 1 µL/mL benzonase (Sigma-Aldrich, E1014) and 10 µL/mL Triton X-100 (Sigma-Aldrich, X100) and 5 mg/mL type I Collagenase (Worthington) at 4°C for 48 hours, followed by centrifugation for 15 min at 4255×g and incubation with additional benzonase. Following a second centrifugation at 5000×g, cell-free supernatant containing virus was aliquoted for single use and long-term storage at-80°C. For viral genome equivalents (VGE) quantification, 10 μL of virus stocks were treated with equal volume viral lysis buffer (0.1% Proteinase K, 0.5% SDS, 25 mM EDTA) at 55 °C for 30 min, and total DNA amount were estimated by agarose gel electrophoresis and comparison to 2 μL Quick-Load Purple 1 kb Ladder (NEB, N0552). Then VGE was converted as 0.82 ng DNA = 10^8^ VGE.

### Primary keratinocyte infection studies

NIH-3T3 murine fibroblasts (ATCC, CRL-1658) were grown in Dulbecco’s modified Eagle medium (DMEM, Invitrogen, 11960–069) supplemented with 5% fetal bovine serum (ThermoFisher Scientific, A4736401). Early-passage murine keratinocytes on FVB/N background were isolated and cultured as described (24, 25). In brief, neonates were euthanized and whole skin was physically disassociated, immersed in 0.25% trypsin overnight at 4°C. Epidermis was separated from dermis and prepared as single cell suspension using gentleMACS Tissue Dissociators (Miltenyi Biotec), seeded in mitomycin-treated NIH-3T3 cells (feeder layer). Murine keratinocytes were cultured in F-medium [3:1 F12 (Invitrogen, 11765–062)/DMEM (Invitrogen, 11960–069), supplied with 5% Fetal bovine serum (ThermoFisher Scientific, A4736401), 2 mM L-Glutamine (Invitrogen, 25030081), 0.4 μg/mL hydrocortisone (Sigma-Aldrich, H4001), 8.4 ng/mL cholera toxin (Calbioche, 227036), 10 ng/mL EGF (Invitrogen, PHG0311), 24 μg/mL adenine (Sigma-Aldrich, A-2786), 5 μg/mL insulin (Gemini, 700–112P), 100 U/mL Penicillin-Streptomycin (ThermoFisher Scientific, 15140163), 10 µM dihydrochloride (MedChemExpress, HY-10583)]. 100,000 cells were seeded in 6-well plate 1 hour prior to infection without feeder layer. Virus stock was diluted in PBS with a 10-fold serial dilution from 10^4^ to 10^8^ VGE. Mock infection was prepared as equal volume of PBS. Mock or viral prep were then mixed with culture media and added to cell cultures. RNA was harvested 48 hours or 96 hours post infection (p.i.). For viral replication experiments, 100,000 cells were seeded without feeder layer, infected with 10^8^ VGE virus. Cells were passaged at day 2 and every 4 days until day 14, and 4/5 of total cells were harvested for DNA extraction. The remaining cells were passaged by removing feeder layer and seeding in a new feeder layer-coated 6 cm dish.

### RNA/DNA extraction and (RT-)qPCR

Genomic DNA and total RNA were extracted using the DNeasy Blood and Tissue Kit (Qiagen, 69504) or RNeasy Mini kit (Qiagen, 74104), respectively, following the user manual. Extracted DNA or RNA was quantified by Nanodrop and normalized between samples. cDNA was synthesized using QuantiTect reverse transcription kit (Qiagen, 205311). Quantitative (q)PCR on DNA or cDNA samples was performed using an ABI Prism 7900HT sequence detection system (Applied Biosystems) or LifeCycler 480 II instrument (Roche). RT-qPCR primers are listed in **Table S1**.

### Immunoblotting

Cells were harvested and resuspended in HED buffer [25 mM Hepes, 5 mM EDTA, 10% glycerol, 1 mM DTT, and 1× protease inhibitor cocktail tablet (cOmplete, Roche, 04693132001)] and underwent multiple freeze and thaw cycles followed by 20 min water sonication. Proteins were then collected from supernatant of lysates, quantified by Bradford assay (ThermoFisher Scientific, 23200), fractionated using a 4-20% gradient SDS-PAGE gel (Bio-Rad, 5671094), and then transferred to a polyvinylidene difluoride Immobilon-FL membrane (Sigma-Aldrich, IPFL00005). Membranes were washed in PBS containing 0.1% Tween-20 (PBST, Alfa Aesar, J20605-AP) and 1X casein buffer (Sigma-Aldrich, B6429) for blocking non-specific binding. The membranes were incubated with a primary rabbit α-human A3A/B/G mAb [1:1000 (26)] and a mouse α-actin (1:5000, Sigma-Aldrich, A1978) or mouse α-tubulin (1:10000, Sigma-Aldrich, T5168) at 4°C overnight. Membranes were then washed in PBST and incubated for 1 hour with secondary α-rabbit HRP (1:10000, Cell Signaling Technology, 7074) and goat α-mouse 800 (1:10000, LI-COR, 926-32210). These membranes were then washed 5 times in PBST and one time in PBS for 5 min each, then imaged using an Odyssey M imager (LI-COR). SuperSignal West Femto Maximum Sensitivity Substrate (ThermoFisher Scientific, 34096) was applied to membrane 5 min prior to imaging to detect HRP.

### DNA deamination activity assays

Whole cell extracts (WCE) in 100,000 cells/100 µL HED buffer were collected and quantified as above, 75 µg total protein was then co-incubated with 800 nM 5′-ATTATTATTATT<u>C</u>GAATGGATTTATTTATTTATTTATTTATTT-FAM oligo at 37°C with 1 µg RNase A (Sigma-Aldrich, R5503)and 0.1 U uracil DNA glycosylase (New England Biolabs, NEB, M0280) for 24 hours. Equal volume of HED buffer co-incubated with oligo was used as negative control, and recombinant APOBEC3A (A3A) protein co-incubation was used as positive control. Samples were heated to 98°C for 10 min supplemented with 100 mM NaOH to induce abasic site cleavage. Samples were separated by 15% TBE-Urea PAGE gel electrophoresis and imaged using an Odyssey Classic scanner or an Odyssey M imager (LI-COR).

### Plasmids

pRSH11141=pcDNA3.1-MmuPV1 or pRSH9977=pcDNA3.1-MND-eGFP (hereafter as pMND-eGFP) were transformed into competent 10-beta *E. coli* (NEB, C3019H), cultured in LB, and extracted using PureLink HiPure Plasmid Maxiprep Kit (Invitrogen, K210007). Virus genomes were prepared by recircularization (hereafter as pMmuPV1) of pRSH11344=pcDNA3.1-MmuPV1 as described (27, 28). Briefly, 50 μg pRSH11344=pcDNA3.1-MmuPV1 was incubated with 50U BamHI-HF (NEB, R3136) at 37°C overnight and then purified with GeneJET Gel Extraction and DNA Cleanup Micro Kit (ThermoFisher Scientific, K0831). Linearized plasmid DNA was then incubated in 6 mL ligation buffer with 10U T4 DNA ligase (NEB, M0202) at 16°C overnight. Recircularized genomes were then precipitated with GeneJET Gel Extraction and DNA Cleanup Micro Kit.

### Transfection assays

2.5*10^4^ cells were seeded in Nunc Lab-Tek II Chamber Slide System (ThermoFisher Scientific, 154534PK) for immunofluorescence (IF) staining. A total of either 500 ng pMND-eGFP or 250 ng pMND-eGFP plus 250ng pMmuPV1 were transfected into each well using Lipofectamine 3000 (Invitrogen, L3000001) the next day. Media were changed 24 hours later, and cells were fixed 48 hours post-transfection. For the deamination activity assay, 2*10^5^ cells were seeded in a 6-well plate, and a total of 1000 ng pMND-eGFP or 1000 ng pMmuPV1 was transfected as aforementioned. Cells were harvested and proteins were extracted 48 hours post-transfection using HED buffer.

### Immunofluorescent microscopy

Cells were fixed in 4% paraformaldehyde in PBS for 15 min at room temperature. Cells were washed with cold PBS for 5 min and then permeabilized in 1% Triton X-100 in PBS for 10 min at 4°C. Cells were washed once with cold PBS for 5 min. Cells were incubated in a blocking solution of 5% normal goat serum (Gibco, 16210064) and 1% Triton X-100 in PBS for 1 hour at room temperature on a rocker. Primary antibody in blocking solution was added, and the cells were incubated overnight at 4°C. Cells were washed with PBS 3× for 5 min at room temperature. Cells were incubated with secondary antibodies at a concentration of 1:1000 in blocking solution for 2 hours in the dark at room temperature with gentle rocking. Cells were washed with PBS at room temperature before being imaged on a Nikon Inverted TI-E Deconvolution microscope (Nikon).

Cells were counterstained with Hoechst 33342 Ready Flow™ Reagent (Invitrogen, R37165) to visualize DNA content. The primary antibody was a custom rabbit α-human A3A/B/G mAb 1:150. Secondary antibody was goat anti-rabbit IgG (H + L) Highly Cross-Adsorbed Secondary Antibody, Alexa Fluor™ 594 (Invitrogen, A-11037).

### Histology and immunohistochemistry staining

Murine ears were bisected and fixed in 4% paraformaldehyde in PBS for 24-48 hours at 4°C. Fixed tissues were then transferred to 70% ethanol, processed, embedded in paraffin and sectioned using a manual rotary microtome (5 μm sections; Leica RM2235). Every 10th section was stained with hematoxylin and eosin (H&E). Immunohistochemistry staining was performed as previously described (22, 29, 30). Briefly, slides were deparaffinized at 65°C, and rehydrated with CitriSolv (Decon Labs, 1601) and graded alcohols. The slides were immersed into 1X Reveal Decloaker (BioCare Medical, RV1000M) water bath at 95°C for epitope retrieval, then soaked in 3% H_2_O_2_ diluted in TBST to suppress endogenous peroxidase activity. Background Sniper (BioCare Medical, BS966) was used to block nonspecific binding, followed by primary antibody incubation overnight at 4°C. Then slides were incubated with Novolink Polymer (Leica Biosystems, RE7280), covered with the Novolink DAB substrate kit (Leica Biosystems, RE7280), and counterstained with Mayer’s hematoxylin solution (Electron Microscopy Sciences, 26252-01). Finally, slides were dehydrated and cover-slipped with Permount mounting media (ThermoFisher Scientific, SP15100). Primary antibodies used for detection were the anti-A3B/A/G mAb 5210-87-13 [1:350, (26)], an anti-MmuPV1 E4 pAb [1:30000, kind gift from Dr. Doorbar Lab, Cambridge University, (31)], and an anti-Ki67 mAb (1:500, Invitrogen, MA5-14520). QuPath software (version 0.6.0.) was used to detect and analyze A3B, E4, and Ki67 positive cells within cutaneous dysplastic lesions.

### Pathology

All six H&E-stained slides from each ear were independently evaluated by a pathologist (P. P. A.) blinded to the genotypes. Pathological changes were assessed and scored as non-dysplastic (ND), low-grade dysplasia (LGD), high-grade dysplasia (HGD), or carcinoma *in situ* (CIS). HGD was defined as cytologic and architectural epithelial aberrations extending through the basilar two-thirds of the epidermis and sparing the superficial keratin layer, whereas CIS occupied the entire epidermal thickness. Conversely, LGD was defined as maturational disorganization limited to the basilar one-third to one-half of the epidermis. LGD was ultimately excluded from the analysis owing to the, overall, attenuated thickness of the ear epidermis. The H&E-stained tissue section displaying the most severe microscopic features, i.e., degree of squamous dysplasia, was utilized for the final histopathologic assessment for each ear specimen.

### Differential DNA denaturation PCR (3D-PCR)

Genomic DNA was extracted as described above. 25ng of total DNA was added into 2X Apex Taq RED Master Mix (Genesee Scientific, 42-138) with E2 or Locus Control Region (LCR) primers listed in **Table S1**. Equal amount of pMmuPV1 was used as negative control, and pMmuPV1 pre-incubated with recombinant A3A at 37°C overnight was used as positive control. PCR reactions were performed using Mastercycler X50s (Eppendorf) with a gradient denaturation temperature for 30 sec, annealing at 60°C for 30 sec, and extension at 72°C for 1 min for a total of 32 cycles. Then PCR products were separated by 2% agarose gel and imaged by Gel Doc XR+ Gel Documentation System (Bio-Rad, 170-8170).

### MmuPV1 sequencing

Whole genome of MmuPV1 was amplified using Phusion High-Fidelity PCR Master Mix (NEB, M0531) with primers listed in **Table S1**. PCR reactions were performed using Mastercycler X50s (Eppendorf) with 98°C for 30 sec, annealing at 60°C for 30 sec, and extension at 72°C for 5 mins for a total of 30 cycles. Amplicons were purified with GeneJET Gel Extraction and DNA Cleanup Micro Kit, and inserted into pJet plasmid using CloneJET PCR Cloning Kit (ThermoFisher Scientific, K1231), then transformed into competent 10-beta *E. coli*. (NEB, C3020K). Individual clones were picked and cultured and plasmid DNA was extracted using QIAprep Spin Miniprep Kit (Qiagen, 27104). Plasmid DNA from each clone was sent to Eurofins Scientific for whole plasmid sequencing. The sequences were aligned back to pMmuPV1 sequence using SnapGene (v6.0.2).

### Statistical analyses and visual presentation

Mann-Whitney U test was used to compare median values between two groups as they had non-normal distributions. Kruskal-Wallis test was used to compare values between three groups as they had non-normal distribution, followed by post hoc pairwise comparisons with Bonferroni correction. Fisher’s Exact test was used to compare two independent categorical groups. Data were analyzed and visualized using R version 4.2.2. All figures were created using Illustrator 2026 (Adobe), and illustrations in **Fig. 1B** and **Fig. 3A** were made with BioRender [Harris, R. (2026) https://BioRender.com/9l7j6im].

**FIG 1.**
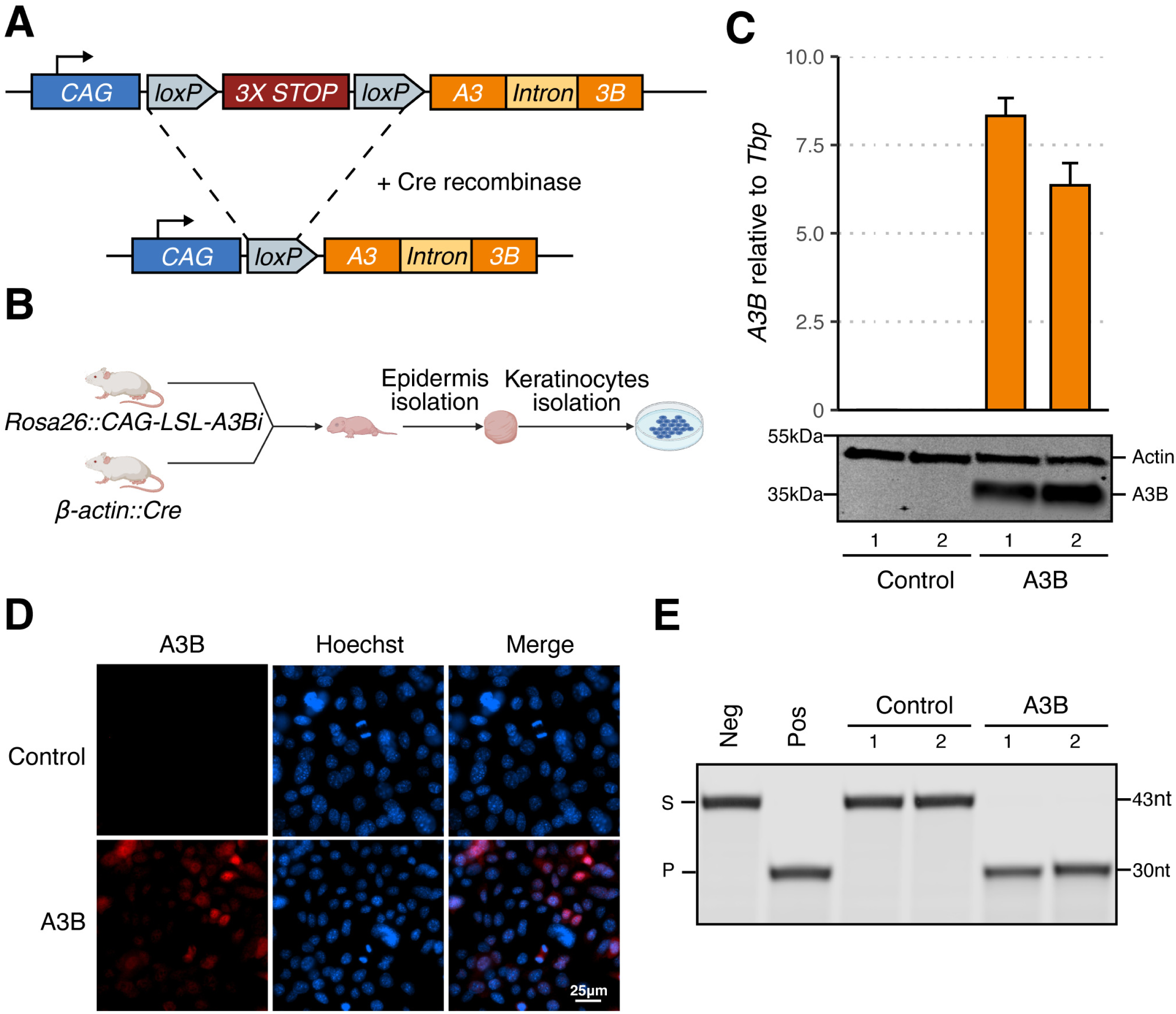
Establishment of an inducible human A3B mouse model. (A) Schematic of human *A3B* minigene integrated into the murine *Rosa26* locus before and after Cre-mediated recombination. (B) Workflow for establishing primary murine keratinocyte cultures. (C) RT-qPCR and immunoblot results for two control (rep1 MKB051, rep2 MKB052) and two A3B-expressing keratinocyte cultures (rep1 MKB056, rep2 MKB048). (D) IF microscopy of control (rep1) and A3B (rep1) murine keratinocytes (blue: Hoechst; red: A3B). (E) DNA deamination activity of extracts from the indicated cultures (S: substrate; P: product). The negative control is substrate incubated with HED buffer, and the positive control is substrate incubated with HED buffer containing recombinant A3A.

## RESULTS

### Engineering human A3B expression in FVB mice

A human *A3B* minigene was engineered into the non-essential *Rosa26* locus of FVB/N mice by homologous recombination in individual embryos, followed by founder screening and germline transmission validation (**Methods**). This minigene has a *CAG* promoter, a triple transcription termination cassette flanked by two *loxP* sites, and an intron-containing *A3B* open reading frame (**Fig. 1A**). Human A3B expression occurs upon Cre-mediated recombination to remove the triple stop cassette. After establishment of the initial *Rosa26::CAG-LSL-A3Bi* line, representative animals were crossed with *β-actin::Cre* mice to yield pups expressing human A3B in all tissues (*i.e*., full body). These animals appeared healthy apart from male-specific infertility as reported recently for the same minigene construct in a C57BL/6J background (22).

Neonatal animals were used to establish primary keratinocyte cultures expressing human A3B (*Rosa26::CAG-L-A3Bi*; *β-actin::Cre*) or not expressing A3B as controls (wildtype animals or unrecombined *Rosa26::CAG-LSL-A3Bi* animals). Two A3B-expressing primary keratinocyte cultures (hereafter designated the A3B group; rep1 MKB056, rep2 MKB048) and two littermate control cultures (the control group; rep1 MKB051, rep2 MKB052) were established from the same litter (**Fig. 1B**). As expected, A3B was expressed at both the RNA and protein levels in A3B cultures but not in control cultures (**Fig. 1C**). No change in morphology or stratification was observed in A3B expressing cells compared to controls (data not shown). A3B protein expression was also confirmed by direct IF microscopy and, after preparation of cell extracts, by single-stranded DNA C-to-U deamination activity assays (**Fig. 1D-E**). Moreover, as in human cells, A3B localizes primarily to the nuclear compartment in primary murine keratinocytes (**Fig. 1D**).

### MmuPV1 is resistant to restriction by human A3B in primary murine keratinocytes

MmuPV1 virus stocks were prepared in immunodeficient mice as reported (23). To ask whether human A3B can restrict MmuPV1, 100,000 A3B-expressing or control primary murine keratinocytes were infected with a log_10_ gradient of MmuPV1 viral genome equivalents (10^4^ to 10^8^ VGE), which helped to ensure low (<1) to high (>>1) multiplicity of infection (MOI; experimental workflow in **Fig. 2A**). MmuPV1 E1^E4 transcript levels were quantified by RT-qPCR and, surprisingly, A3B expression had no negative effect at 5 different viral MOIs, either 48 or 96 hours p.i. (**Fig. 2B**). If anything, A3B triggered a slight increase in viral transcript levels at several of the time points post-infection.

**FIG 2.**
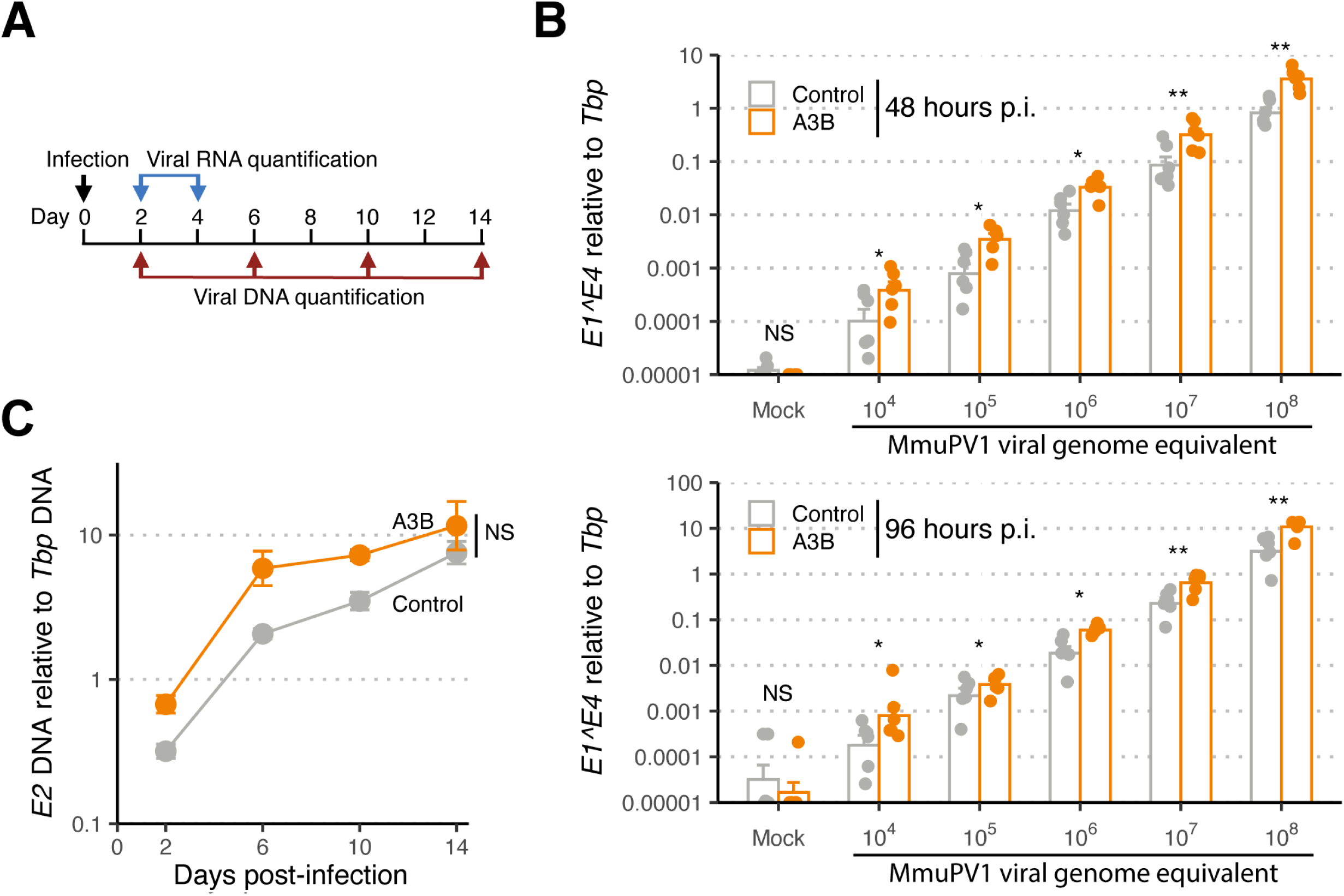
MmuPV1 is resistant to A3B restriction *in cellulo*. (A) Schematic of viral Infection assay and timepoints for RNA and DNA quantification. (B) RT-qPCR results for MmuPV1 E1^E4 spliced transcript normalized to murine *TATA binding protein* (*Tbp*) mRNA transcript 48 hours or 96 hours p.i. (top and bottom bar graphs). Data from two control and two A3B groups were combined [n = 3 infections per group with individual data points and means shown; *, *p* < 0.05; **, *p* < 0.01; ***, *p* < 0.001; and *p* ≥ 0.05, non-significant (NS) by Mann-Whitney U test]. (C) qPCR results for MmuPV1 E2 DNA relative to *Tbp* genomic DNA (mean +/-SD from n = 2 independent experiments, each with 2 cultures; *p* = 0.11 by two-factor, two-way ANOVA).

**FIG 3.**
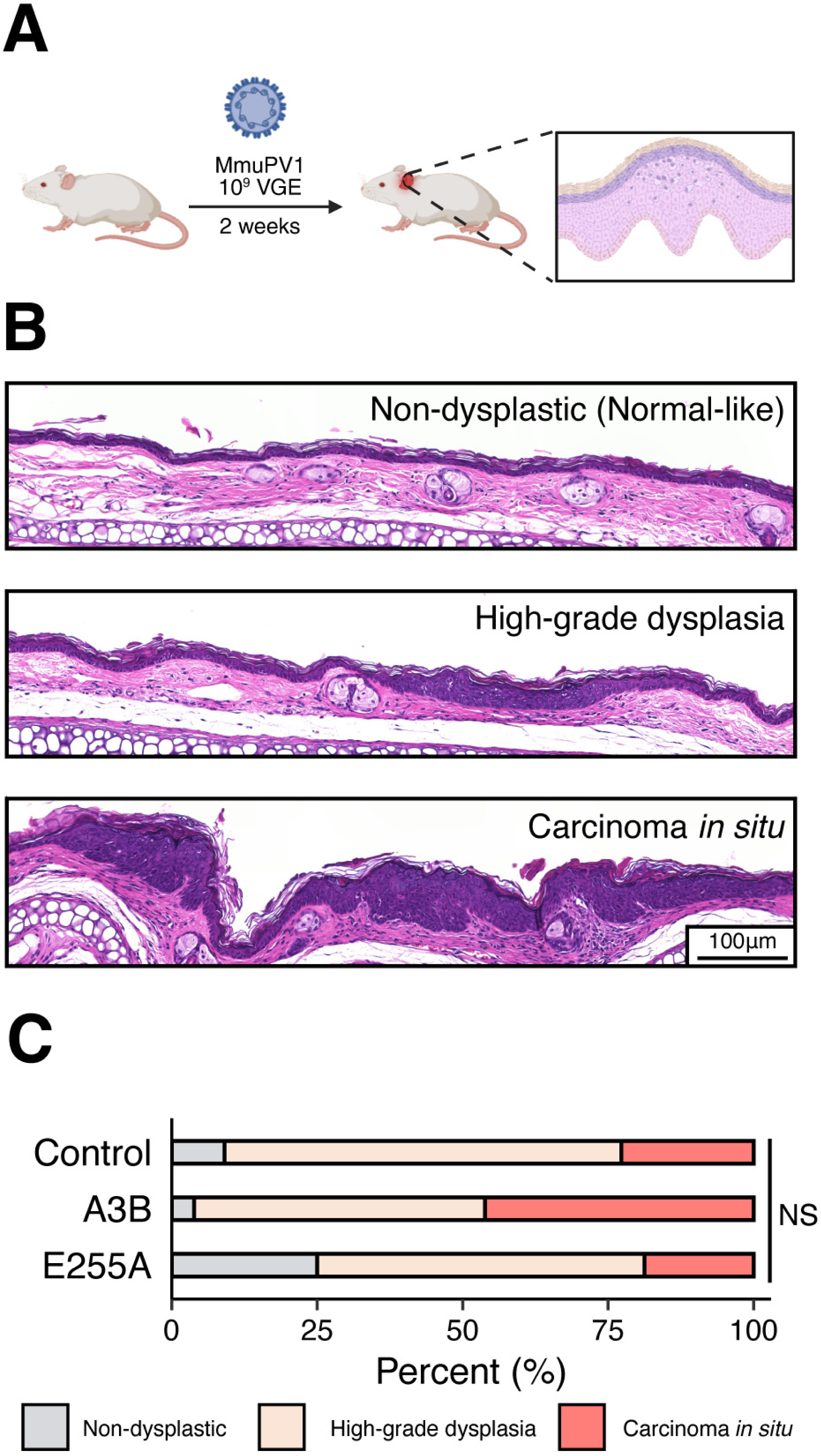
MmuPV1 is resistant to A3B restriction *in vivo*. (A) Workflow for the MmuPV1 early-stage ear infection model. (B) Representative H&E photomicrographs of the histopathologic spectrum of ear skin lesions including non-dysplastic (normal-like), high-grade dysplasia, and carcinoma *in situ*. (C) Summary of overall histopathologic grading of cutaneous dysplastic lesions of the ear between control (n = 22), A3B (n = 26), and E255A (n = 16) groups of infected animals (NS, not significant; *p* = 0.40 by Kruskal-Wallis test including all three groups, and *p* > 0.05 by Fisher’s Exact test for all pairwise combinations).

We next asked whether human A3B expression affects MmuPV1 DNA replication and maintenance. This was done by infecting cells with 10^8^ VGE and then using qPCR to quantify MmuPV1 E2 DNA accumulation over a 14-day period (experimental workflow in **Fig. 2A**). As above for viral mRNA levels, viral genomic DNA copies remained comparable between A3B and control groups (**Fig. 2C**). Moreover, if anything, viral DNA levels increased slightly more rapidly in A3B-expressing primary murine keratinocytes. Similar results were obtained with 2 independent A3B-expressing primary murine keratinocyte cultures (rep3 MKB067, rep4 MKB069) and 2 independent control cell cultures (rep3 MKB070, rep4 MKB071) derived from the progeny of a different pair of parent animals (**Fig. S1-2**). Collectively, these RT-qPCR and qPCR results combine to indicate that MmuPV1 is somehow capable of resisting the antiviral activity of human A3B.

### MmuPV1 also resists human A3B *in vivo* in an early-stage ear infection model

Similar to HPV, MmuPV1 requires a fully stratified keratinocyte epithelium to complete its life cycle (32–34). However, early-passage murine keratinocytes in monolayer culture remain basal-like and undifferentiated, limiting their ability to model the complete viral life cycle. To overcome this limitation, we turned to an *in vivo* cutaneous MmuPV1 infection model, enabling us to determine whether *A3B* restricts any stage of the viral life cycle including late, differentiation-dependent stages (35). In brief, the inner side of both ears of each animal was scraped with a needle, and a total of 10^9^ VGE was applied to wound sites. Two weeks later, animals were sacrificed, and ear tissues harvested for histopathologic examination (workflow in **Fig. 3A**). In addition to the A3B and control F1 animals described above, these experiments also included an additional group of F1 animals that expresses a catalytically defective A3B protein (*i.e*., doubly heterozygous animals derived from crossing *Rosa26::CAG-LSL-A3Bi-E255A* and *β-actin::Cre* mice and hereafter called the E255A group).

Epithelial alterations were assessed and scored as non-dysplastic (normal-like), high-grade dysplasia, and carcinoma *in situ* (representative images in **Fig. 3B**). Under these infection conditions, most infected ear tissues exhibited high-grade dysplasia or carcinoma *in situ*. However, despite clear virus-dependent phenotypes, no histopathologic differences were observed that could be attributable to A3B between these three groups (**Fig. 3C**).

To further investigate these infected tissues and provide additional controls, we asked whether A3B is expressed in the same cells as virus infection. This was done by staining serial sections of each tissue with an anti-A3B mAb (26), an anti-MmuPV1 E4 pAb (31), and an anti-Ki67 mAb. As expected, A3B protein is expressed in the nuclei of whole skin layers in A3B and E255A group mice but not in control mice (**Fig. 4A-B**). A comparable level of E4 immunopositivity was observed in cutaneous dysplastic lesions of the ear between control and A3B or E255A group ears, suggesting MmuPV1 is still present in stratified skin layers in most dysplastic lesions regardless of A3B expression (**Fig. 4A-B**). Within most dysplastic lesions, the proliferative marker Ki67 highlighted predominantly basal and suprabasal cells, with no statistically significant differences observed regarding Ki67 expression levels among the three groups (**Fig. 4A-B**). Collectively, these histologic and immunohistochemical findings indicate that MmuPV1 is resistant to A3B’s restriction activity through the entire viral life cycle.

**FIG 4.**
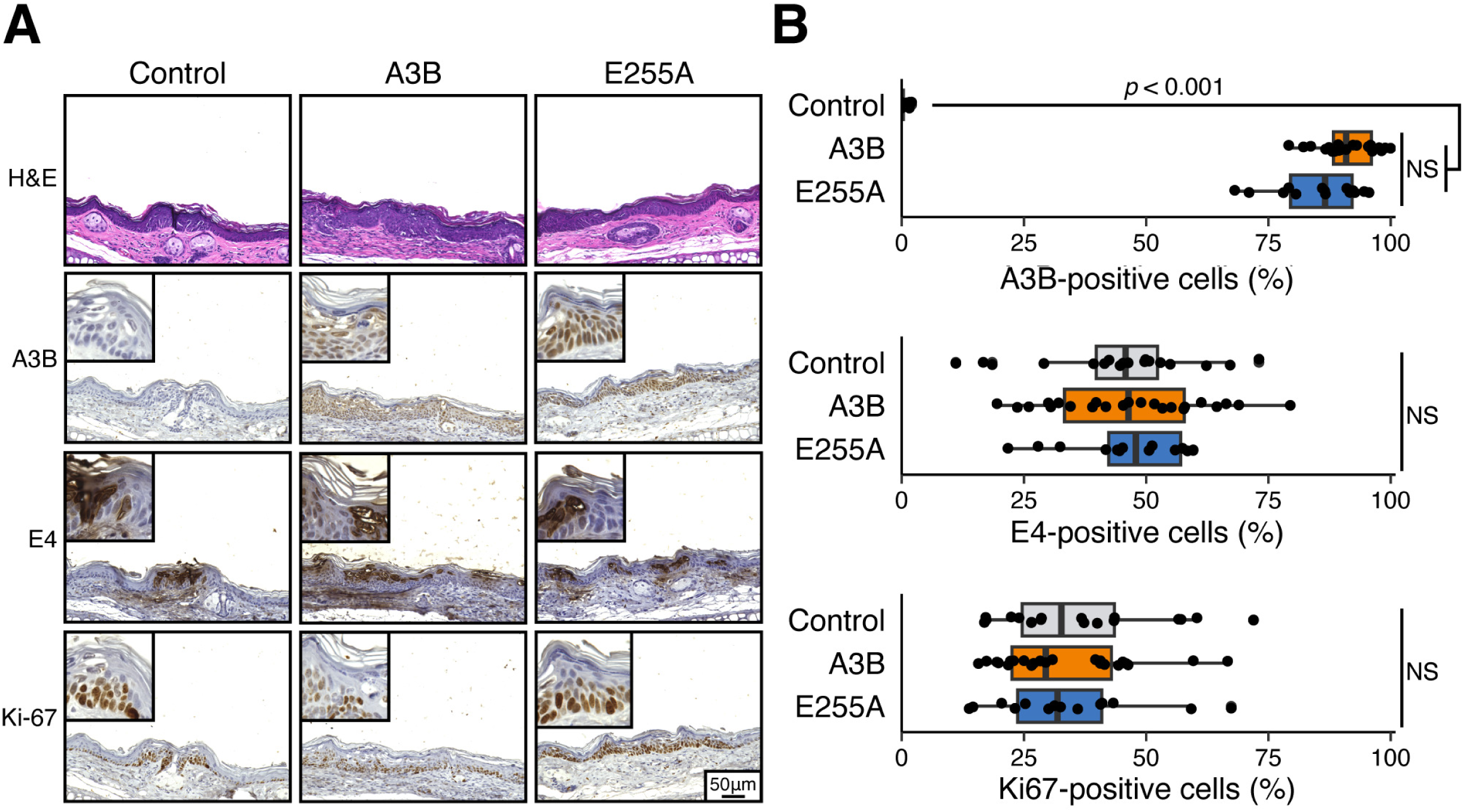
MmuPV1 is proliferative in A3B-expressing cells. (A) Histopathologic and immunohistochemical (IHC) characteristics of MmuPV1-infected ear dysplasias. Representative medium (20X objective) and high-power (40X objective, top left) images are shown. (B) IHC quantification of A3B, MmuPV1 E4, and Ki67 in each of the infected groups (E4 negative cases were excluded from analysis; control, n = 19; A3B, n = 24; E255A, n = 14; statistical comparisons by Kruskal-Wallis test; NS, not significant).

### MmuPV1 appears resistant to A3B mutagenesis

Last, we used differential DNA denaturation PCR (3D-PCR) to ask if human A3B is capable of deaminating MmuPV1 genomic DNA *in vivo*. 3D-PCR preferentially amplifies C-to-U (or C-to-T) hypermutated sequences by virtue of lower denaturation temperatures, in comparison to fully C:G base-paired parental sequences (13, 36, 37). This powerful technique was therefore applied to viral *Locus Control Region* (*LCR*) and *E2* region DNA from day 14 culture DNA extracts. These two loci are essential for virus replication, and mutations in these regions have been correlated with defective replication (38, 39).

Recircularized MmuPV1 genome from plasmid (hereafter pMmuPV1, **Methods**) alone or co-incubated with recombinant A3A protein was used as negative and positive control templates, respectively, for 3D-PCR. These controls clearly showed that viral DNA can be deaminated by an APOBEC enzyme, as evidenced by the appearance of amplicons at much lower denaturation temperatures following treatment with A3A (**Fig. 5A**). However, amplicons from both the A3B and control groups were indistinguishable from the negative control indicating a lack of hypermutated viral DNA substrate (**Fig. 5A** and **Fig. S3**). Only *LCR* and *E2* were investigated using the 3D-PCR approach and in the resolution of this technique might be insufficient to reliably detect rare mutations events. We therefore proceeded to recover near-full viral genomes by PCR (all early genes, *L1*, and partial *L2*), and then cloned and sequenced individual plasmids. A total of 6 amplicons from control rep1 and 10 from A3B rep1 were sequenced. Only a few variants were identified in the A3B rep1 sample and only one of which was a C-to-T transition in an APOBEC-preferred trinucleotide motif (**Fig. S4**).

**FIG 5.**
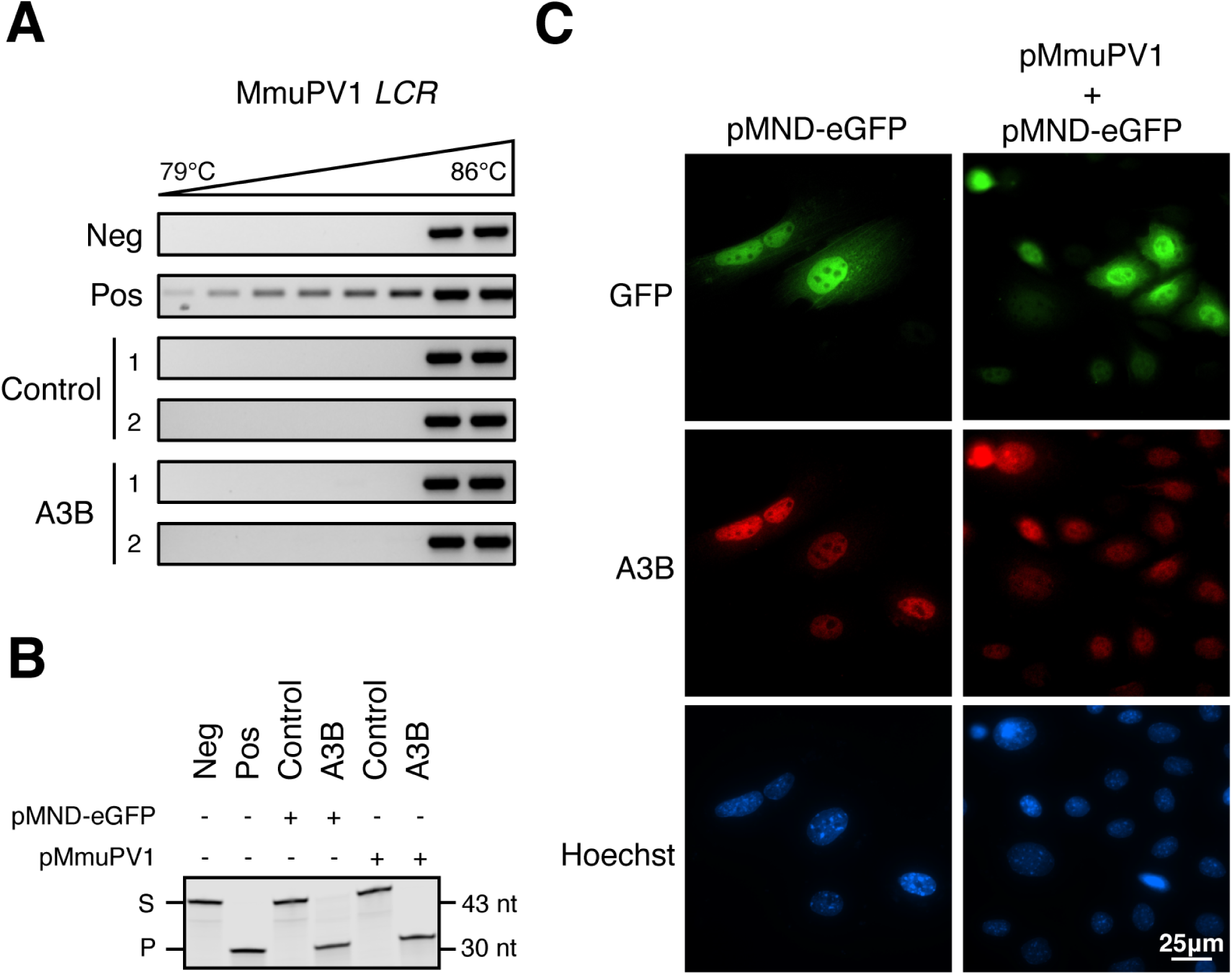
A3B fails to introduce mutations in MmuPV1 genome. (A) Differential DNA denaturation PCR (3D-PCR) of MmuPV1 *LCR* from two control and two A3B-expressing keratinocytes day 14 DNA extracts. Negative control: pMmuPV1; Positive control: pMmuPV1 co-incubated with recombinant A3A. (B) DNA deamination activity of extracts from control rep1 and A3B rep1 protein lysates (S: substrate; P: products). Cells were harvested 48 hours post transfection of either pMND-eGFP or pMmuPV1. Negative control: substrate with HED buffer. Positive control: substrate with HED buffer containing recombinant A3A. (C) Immunofluorescence staining of A3B subcellular localization at 48 hours post-transfection of pMmuPV1 into A3B rep1 cells. pMND-eGFP was included as a co-transfection marker to help identify cells that have likely taken-up pMmuPV1.

To address the molecular mechanism behind the lack of A3B signature mutations in MmuPV1 genomes, we asked if MmuPV1 might somehow attenuate A3B catalytic activity. Murine keratinocyte cultures control rep1 and A3B rep1 were transfected with either pMmuPV1 or pMND-eGFP and harvested 48 hours later for extract preparation and deaminase activity quantification. No differences in A3B deaminase activity were detectable regardless of pMmuPV1 or control pMND-eGFP transfection (**Fig. 5B**). Our previous work demonstrated that Epstein-Barr virus (EBV) and human cytomegalovirus (HCMV) relocalize APOBEC3B (A3B) to the cytoplasm, thereby evading viral genomic DNA mutagenesis by A3B in the nuclear compartment (40, 41). To determine whether MmuPV1 might similarly trigger A3B relocalization, A3B rep1 cell were co-transfected with pMmuPV1 and pMND-eGFP and, 48 hours later, A3B subcellular localization was assessed by IF microscopy. Cell-wide eGFP fluorescence served as a marker for MmuPV1 expression due to a lack of robust antibodies for IF microscopy detection of MmuPV1 proteins. Regardless of eGFP positivity, A3B expression remained predominantly nuclear indicating that MmuPV1 is unable to alter A3B localization (**Fig. 5C**). This result is consistent with A3B nuclear positivity in virus infected murine ear epithelial dysplasias (**Fig. 4A-B**). Collectively, these results indicate that MmuPV1 is not altering A3B activity or subcellular localization.

## DISCUSSION

An absence of clear evidence for HPV restriction by human A3B (**Introduction**) suggests that papillomaviruses may have evolved a counter-defense mechanism. To test this hypothesis, we developed a murine model for inducible expression of human A3B and asked whether this potent antiviral factor is able to restrict murine papillomavirus MmuPV1. A similar cross-species approach was used originally to uncover the retrovirus restriction activity human APOBEC3G (42, 43). However, contrary to expectations, we found that MmuPV1 transcription and replication are unaffected in human A3B-expressing primary murine keratinocytes. We also observed comparable MmuPV1 infection pathologies in an ear infection model regardless of human A3B expression. Furthermore, MmuPV1 genomes from day 14 infected keratinocytes were devoid of APOBEC signature hypermutations despite strong A3B ssDNA deaminase activity and clear nuclear localization. These results combine to indicate that MmuPV1 is somehow able to resist the antiviral activity of human A3B and, by extension, suggest that HPV might have a similarly airtight APOBEC evasion mechanism.

We postulate that papillomaviruses employ a conserved strategy to evade A3B activity and that, based on results here, it is unlikely to be species-specific. This mechanism is unlikely to involve deaminase inhibition and/or relocalization, as for human A3B by herpesviruses such as EBV, KSHV, and HCMV (9, 40, 44–46), because these human A3B activities are not significantly affected by MmuPV1. This mechanism is also unlikely to involve proteasomal degradation of human A3B, as for degradation of related APOBEC3s including APOBEC3G by lentiviruses such as HIV-1, HIV-2, SIV, and MVV (47–53), because A3B protein levels by immunoblot, IF microscopy, and IHC are unaffected by MmuPV1.

The precise nature of the escape mechanism used by papillomaviruses to prevent restriction by human A3B will require further studies to elucidate. For instance, it could be subtle and passive such as an altered epigenomic state, which may serve to directly shield replicating viral genomic DNA from A3B. Alternatively, viral replication intermediates may be physically protected (indirectly shielded) from A3B. For example, viral E1 and E2 proteins are known to recruit multiple host proteins to viral replication factories for transcription, replication, and localization to host chromatin (54–59). Such protein-rich microenvironments might physically prevent A3B from accessing viral ssDNA during transcription (R-loops) and/or replication. Another possibility is that A3B-induced C-to-U deamination may occur but is quickly reversed by host DNA repair pathways before it compromises viral replication. In support of this possibility, DNA damage repair (DDR) pathways are constitutively activated in HPV infected cells and several DDR factors are recruited to viral replication centers (60–64). Other mechanisms may also be plausible. Regardless of the precise solution to this problem, the relationship between HPV infection and human APOBEC3 enzymes is broadly relevant because HPV-positive tumors exhibit some of the highest APOBEC3 mutation loads of all human cancers (10).

## ACKNOWLEDGEMENTS

We thank Amanda Loke, Renee King, and Kerry Smith (University of Wisconsin-Madison) for assistance and insightful suggestions on *in cellulo* infection studies. We are grateful to John Doorbar, Amy Davies, and Egawa Nagayasu (University of Cambridge) for providing an anti-MmuPV1 E4 antibody and to the Gene Targeting & Transgenic Facility (HHMI Janelia Campus) for help establishing the human A3B knock-in models. Research in the Lambert lab is supported by NCI P01-CA022443, NCI R35-CA210807, as well as an American Hair Research Society Mentorship Grant and a UW Academic Staff Professional Development Grant to AB. Research in the Harris laboratory is supported by NIAID R37-AI064046, NCI P01-CA234228, NCI P50-CA247749, and a Recruitment of Established Investigators Award from the Cancer Prevention and Research Institute of Texas (CPRIT RR220053). RSH is an Investigator of the Howard Hughes Medical Institute and the Ewing Halsell President’s Council Distinguished Chair at University of Texas San Antonio.

## Author Contributions

Xingyu Liu, Conceptualization, Data curation, Formal analysis, Investigation, Methodology, Visualization, Writing – original draft, Writing – review and editing | Andrea Bilger, Data curation, Formal analysis, Investigation, Methodology, Writing – review and editing | Denis Lee, Data curation, Formal analysis, Investigation, Validation, Writing – review and editing | Argyris P. Prokopios, Data curation, Formal analysis, Investigation, Writing – review and editing | Jiarui Chen, Formal analysis, Investigation, Validation, Visualization, Writing – review and editing | Ella Ward-Shaw, Data curation, Formal analysis, Investigation, Methodology, Writing – review and editing | Emilia Barreto Duran, Data curation, Investigation, Methodology, Writing – review and editing | Yu-Hsiu T. Lin, Formal analysis, Investigation, Methodology, Validation, Writing – review and editing | Cameron Durfee, Formal analysis, Investigation, Writing – review and editing | Sang Hyun Chun, Formal analysis, Methodology, Writing – review and editing | Mahmoud Ibrahim, Formal analysis, Methodology, Validation, Writing – review and editing | Joshua Proehl, Investigation, Methodology, Validation, Writing – review and editing | Allen J. York, Methodology, Validation, Writing – review and editing | Paul F. Lambert, Conceptualization, Funding acquisition, Resources, Supervision, Methodology, Writing – review and editing | Reuben S. Harris, Conceptualization, Funding acquisition, Resources, Supervision, Methodology, Writing – original draft, Writing – review and editing

